# Construction of High Content Nanobody Library in Mammalian Cells by Linear-double-stranded DNA Based Strategies

**DOI:** 10.1101/2020.06.10.143479

**Authors:** Yanjie Zhao, Weijun Su, Shuai Li

**Affiliations:** Department of Breast Cancer Pathology and Research Laboratory, Tianjin Medical University Cancer Institute and Hospital, National Clinical Research Center for Cancer; Key Laboratory of Cancer Prevention and Therapy, Tianjin; Tianjin’s Clinical Research Center for Cancer, Tianjin 300060, China; School of Medicine, Nankai University, Tianjin 300071, China

**Keywords:** nanobody, complementary determining region (CDR), AND gate, linear-double-stranded DNA (ldsDNA)

## Abstract

Here, we developed two novel methods to construct high content nanobody library in mammalian cells. For the first method, we employed a **<u>l</u>**inear-double-stranded DNA **<u>b</u>**ased **<u>A</u>**ND-**<u>g</u>**ate (LBAG) strategy. Upstream- and downstream-linear-double-stranded DNAs (ldsDNAs), containing identical overlapping sequences, were co-transfected into cultured cell lines to conduct AND-gate calculation to form intact nanobody expression cassette. For the second method, we generated full-length nanobody expression ldsDNA by *in vitro* ligation of restrict cut up- and down-stream ldsDNAs. Then the ldsDNAs were directly transfected into mammalian cells to express nanobody library. Both methods generated over a million different nanobody sequences as revealed by high-throughput sequencing (HTS).

## Results

Here, we took a high-affinity GFP-binding nanobody cAbGFP4 as the backbone of our nanobody library. Via PCR amplification, we split the nanobody expression cassette into a pair of ldsDNAs with 50 bp identical overlapping sequences, and the CDRs of cAbGFP4 nanobody (CDR1 8AA, CDR2 9AA, CDR3 7AA) were replaced by the same number of NNK degenerate codons (Figure 1A). These ldsDNAs were co-transfected into HEK293T cells to conduct AND-gate calculation and generate the intact nanobody expression cassette (Figure 1A) (1). Our previous study showed that ldsDNAs with identical overlapping sequences could conduct in-cell AND-gate calculation through a homologous recombination (HR) pathway (1). Thus, we name this strategy as **<u>l</u>**dsDNA-**<u>b</u>**ased **<u>A</u>**ND-**<u>g</u>**ate, **<u>h</u>**omologous **<u>r</u>**ecombination (LBAG-HR).

**Figure 1.**
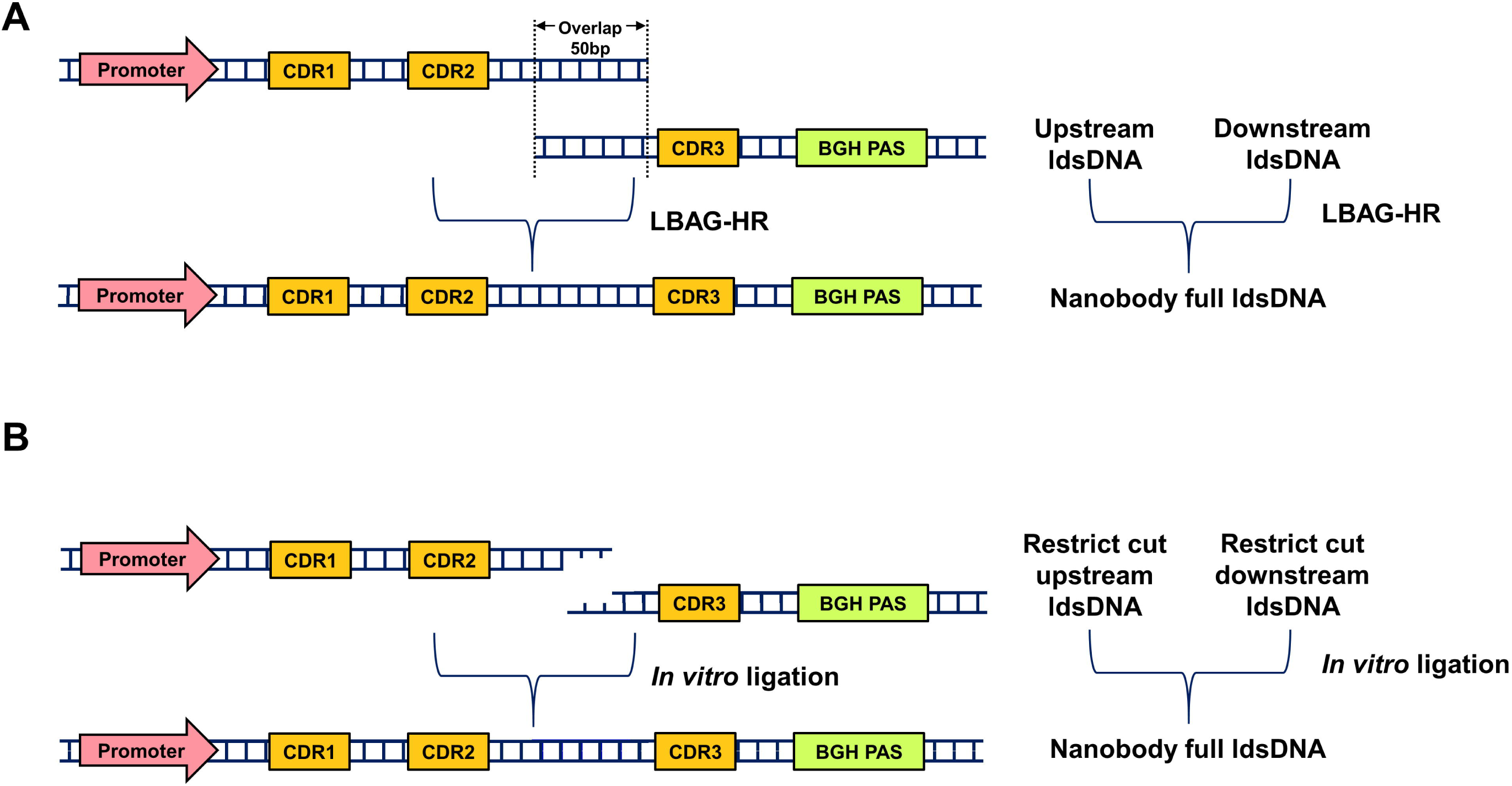
Schematic diagram showing LBAG-HR (A) and *In vitro* ligation (B) nanobody library construction strategies.

We also developed a novel *in vitro* ligation strategy to generate the intact nanobody expression cassette (Figure 1B). Briefly, both the up- and down-stream ldsDNAs were digested by restriction enzyme BssSI to generate sticky ends. Then the restrict-cut ldsDNAs were ligated together by T4 DNA ligase to generate intact nanobody expression ldsDNAs (Figure 1B). At last, these ldsDNAs were transfected into HEK293T cells to express nanobody library. And we name this method as *in vitro* ligation strategy.

Forty-eight hours after ldsDNA transfection, total RNAs were extracted and cDNAs were generated by reverse-transcription. PCR was employed to amplify CDRs of nanobody library. The PCR products were then subjected to high-throughput sequencing (HTS) to explore the characteristics of the library (Table 1). Each strategy contains three biological repeats. LBAG-HR strategy and *In vitro* ligation strategy generated a total of 5,402,539 and 6,322,703 reads, respectively (Table 1). First, we analyzed the length dynamics of the library generated by the two strategies (Figure 2). In LBAG-HR group, a total of 4,305,390 (79.7%) reads show the same length with GFP-nanobody. For *In vitro* ligation group, the full-length reads are 5,182,083 (82.0%) (Table 1). These full-length reads are subjected to the downstream analysis.

**Figure 2.**
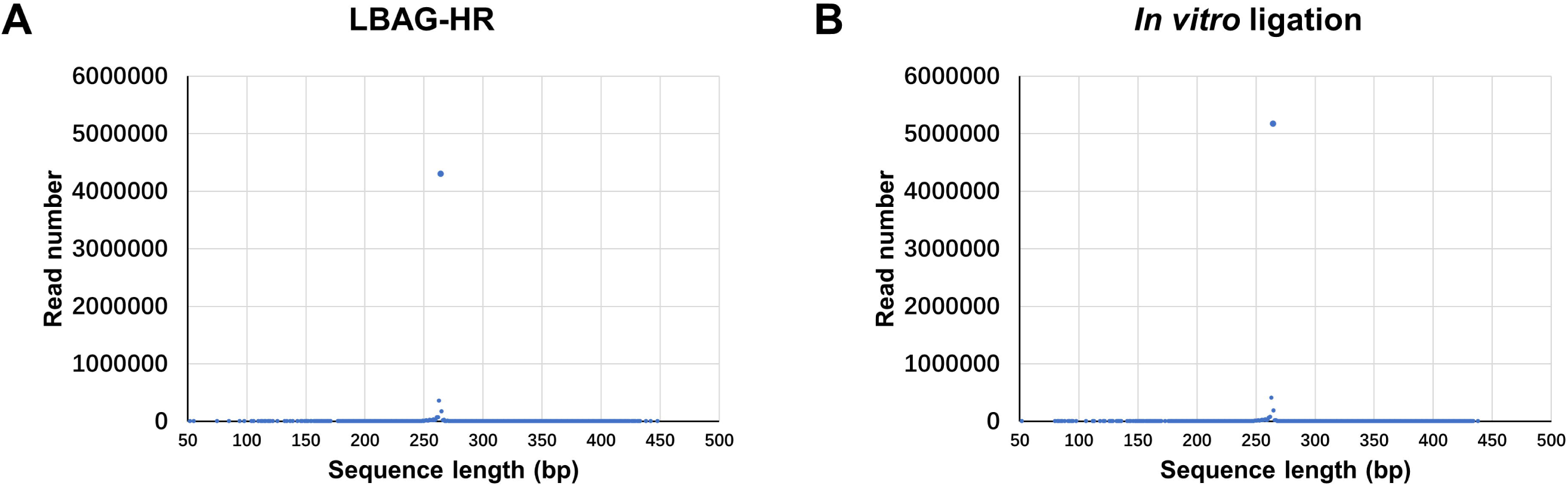
High-throughput sequencing (HTS) revealed the read length distribution of LBAG-HR (A) and *In vitro* ligation (B) nanobody library.

We next analyzed the abundance distribution of full-length reads generated by the two strategies. As shown by Figure 3, about 66% (LBAG-HR) and 64% (*In vitro* ligation) unique sequences have only one reads. In LBAG-HR group, the most abundant sequence has 197 reads. In *In vitro* ligation group, the most abundant sequence has 280 reads. The nucleotide distributions of CDR codons were present in Figure 4.

**Figure 3.**
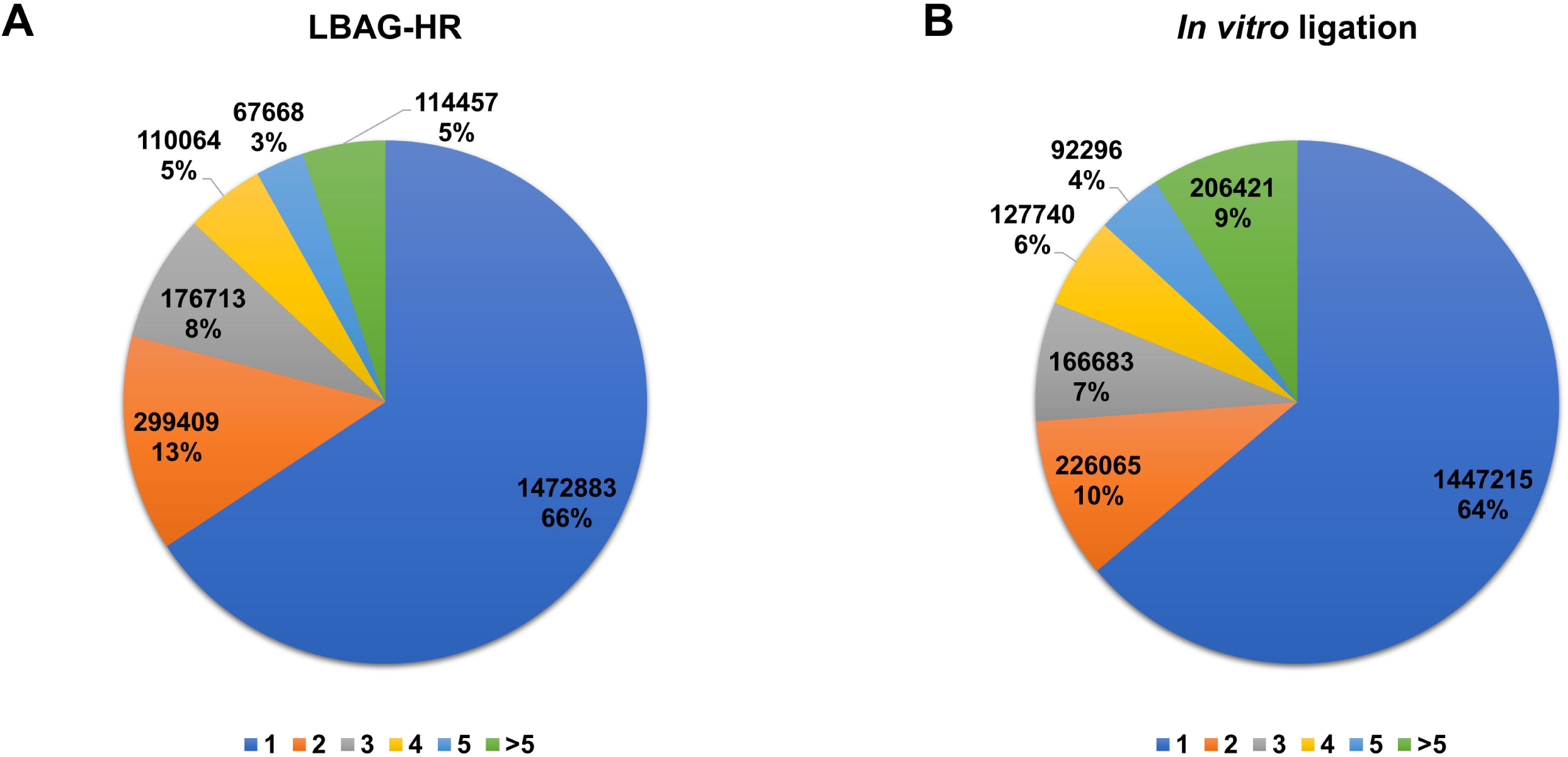
Abundance distribution of full-length nanobody sequences from LBAG-HR (A) and *In vitro* ligation (B) HTS data. Sequence abundance was divided into six groups: 1, 2, 3, 4, 5 and >5.

**Figure 4.**
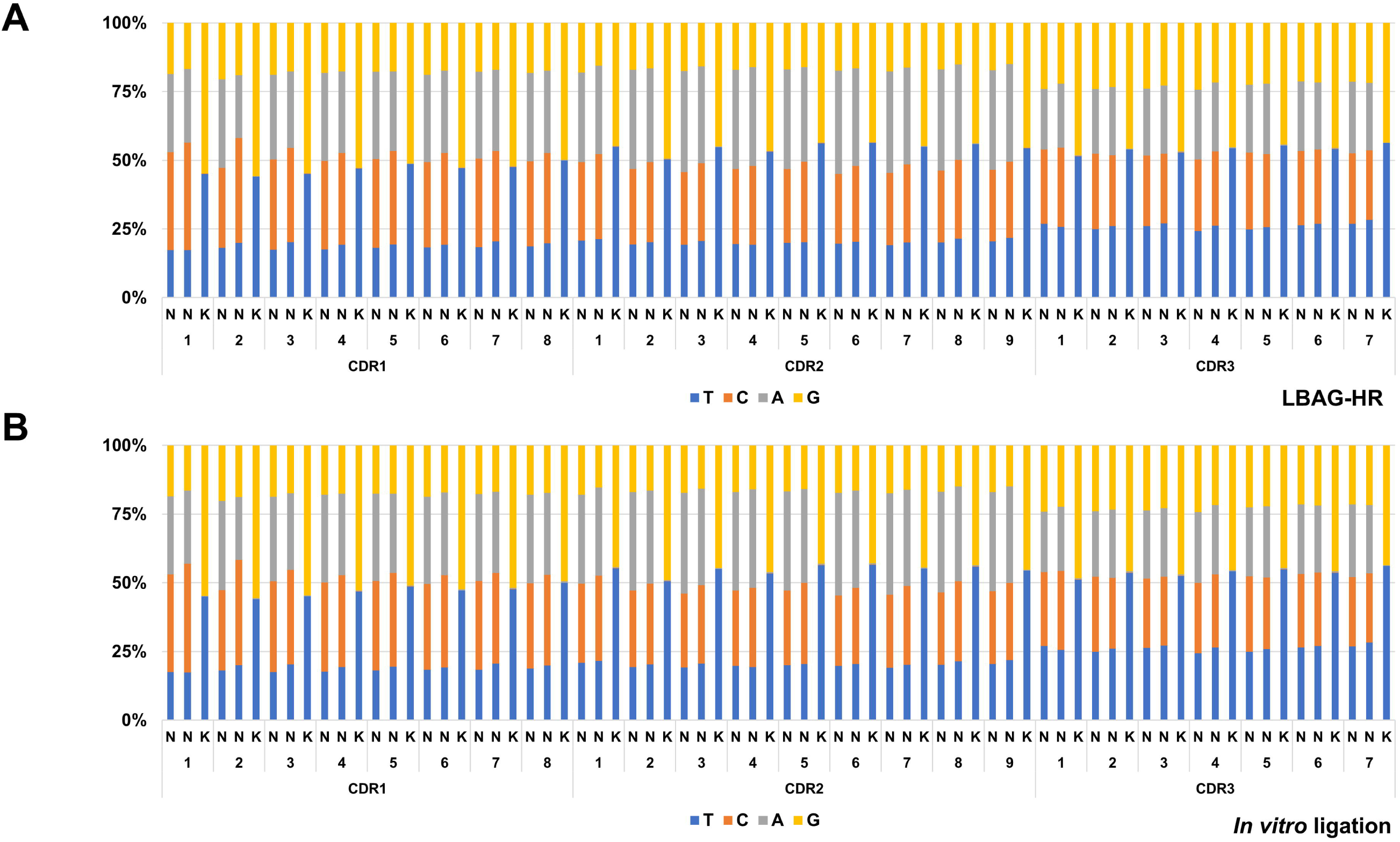
Nucleotide distributions in CDR codons of LBAG-HR (A) and *In vitro* ligation (B) nanobody libraries.

We then translated CDR nucleotide sequences into protein sequences. CDRs’ amino acid compositions were present in Figure 5. Sequences containing pre-stop codon were excluded from further analysis. A total of 1,176,926 and 1,132,956 unique nanobody sequences (in the view of CDRs’ AA sequences) were identified in LBAG-HR group and in *In vitro* ligation group, respectively (Table 1). Venn diagram shows the redundancy of unique nanobody sequences among three repeats of different strategies (Figure 6).

**Figure 5.**
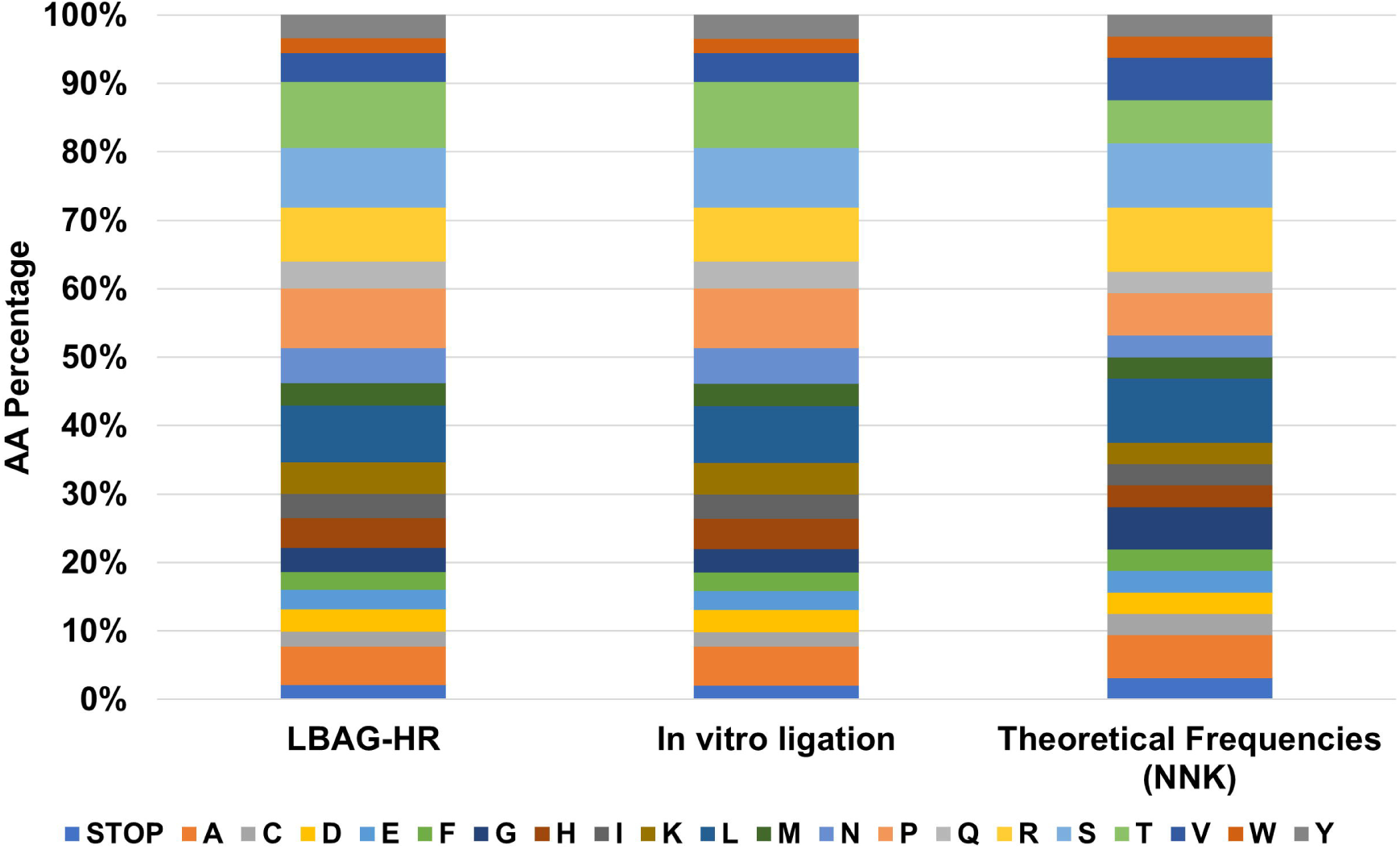
Amino acids composition of CDRs of LBAG-HR and *In vitro* ligation nanobody libraries.

**Figure 6.**
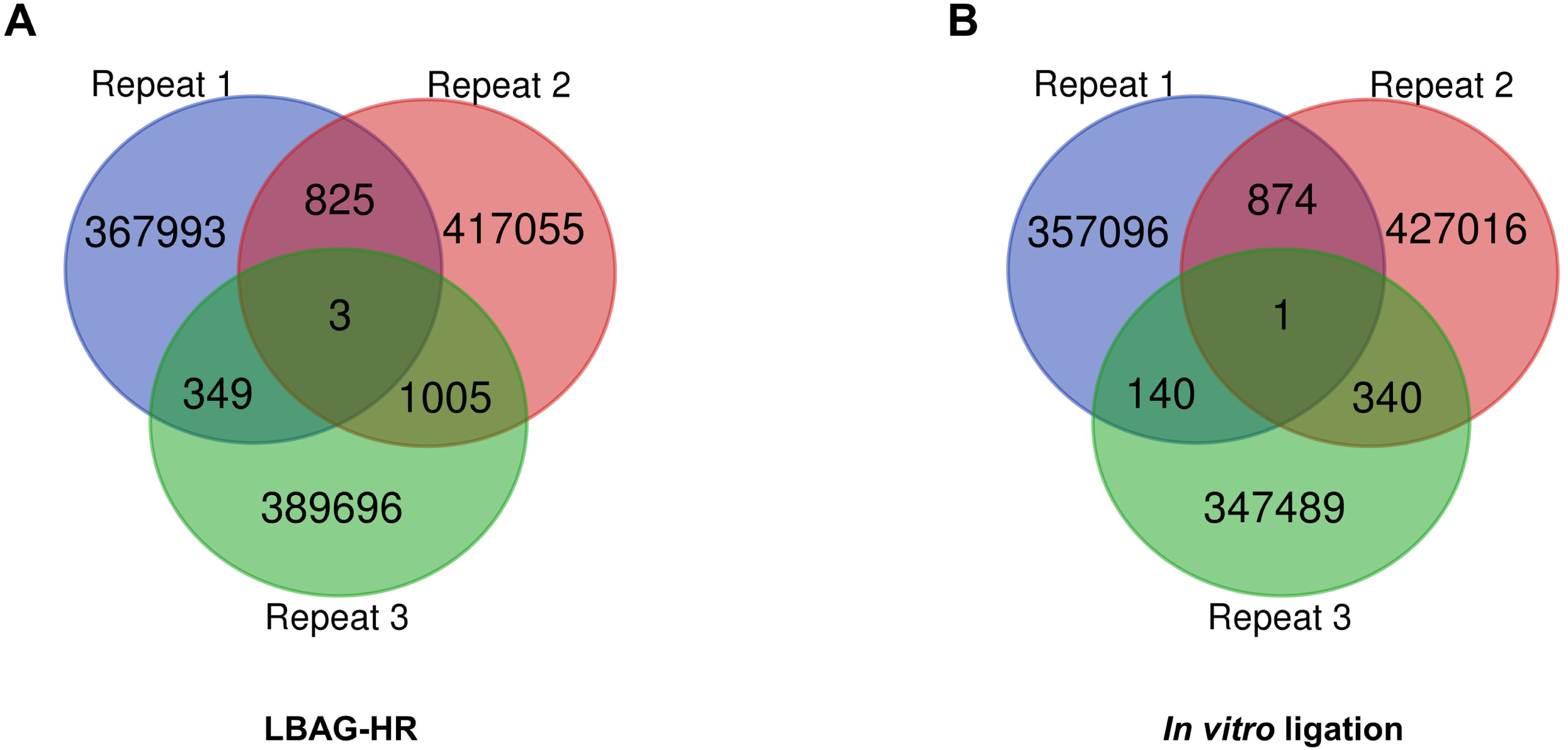
Venn diagram of unique amino acid sequences from three biological repeats of LBAG-HR (A) and *In vitro* ligation (B) nanobody libraries.

## Acknowledgements

This work was supported by the National Natural Science Foundation of China [31870860 to S.L., 31971388 to W.S.]; the State Key Laboratory of Medicinal Chemical Biology (Nankai University) [2018103 to S.L.].

## Competing Interests

The authors have declared that no competing interest exists.

## Notes

### Competing Interest Statement

The authors have declared no competing interest.

