## Supplementary material for "Construction of High Content Nanobody Library in Mammalian Cells by Linear-double-stranded DNA Based Strategies": Table 1

**Table 1.** Characteristics of Nanobody Library.

| **Library Strategy** | **Repeats** | **Reads in Analysis** | **Full Length Nanobody Reads (%)^*^** | **Number of Unique**  **Nanobody Sequence** |
| --- | --- | --- | --- | --- |
| LBAG-HR | Repeat 1 | 1,749,309 | 1,397,756 (79.9) | 369,169 |
|  | Repeat 2 | 1,766,031 | 1,408,849 (79.8) | 418,887 |
|  | Repeat 3 | 1,887,199 | 1,498,785 (79.4) | 391,052 |
|  | **Total^**^** | **5,402,539** | **4,305,390 (79.7)** | **1,176,926** |
| *In vitro* ligation | Repeat 1 | 1,997,936 | 1,639,052 (82.0) | 358,110 |
|  | Repeat 2 | 2,298,395 | 1,872,244 (81.5) | 428,230 |
|  | Repeat 3 | 2,026,372 | 1,670,787 (82.5) | 347,969 |
|  | **Total^**^** | **6,322,703** | **5,182,083 (82.0)** | **1,132,956** |

* % is the percentage of “Full Length Nanobody Reads” in “Read in Analysis”.

** Combination of above three repeats of each strategy.
